## Supplemental information for "*De novo* analysis of bulk RNA-seq data at spatially resolved single-cell resolution"

### Performance evaluation of Bulk2Space

#### Data simulation for the benchmark of Bulk2Space

##### *Collections of single-cell RNA-seq data for data simulation*

To benchmark the performance of Bulk2Space, we designed two types of data simulation procedures. First, the reference data and the synthetic input data were generated from the same dataset, which was termed paired simulation. Ten scRNA-seq datasets were collected for paired simulation, including human and mice primary tissues. Specifically, the 5 human scRNA-seq datasets consisted of the peripheral blood (GSE92495)[1], brain (GSE103723)[2], kidney (GSE121862)[3], liver (GSE124395)[4], and lung (GSE130148)[5]. And the 5 mice scRNA-seq datasets consisted of the brain (GSE60361)[6], kidney (GSE119531)[7], lung (GSE127465)[8], pancreas (GSE84133)[9], and testis (GSE112393)[10] (Fig. S1a). Second, the reference data and the synthetic input data were generated from the same tissue but different datasets, which was termed unpaired simulation. In this simulation, 8 human pancreas scRNA-seq datasets from different resources were collected for unpaired simulation, including a CelSeq (GSE81076)[11], a CelSeq2 (GSE85241)[12], a Fluidigm C1 (GSE86469)[13], a SMART-Seq2 (E-MTAB-5061)[14] and four inDrops (GSE84133)[9] datasets (Fig. S1b). The detailed description of the experimental design for data simulation was summarized in Supplementary Data 1.

##### *Paired and unpaired simulations for evaluation of the deconvolution step of Bulk2Space*

For the deconvolution step, each scRNA-seq data was randomly divided into two parts for paired simulations, with one as the single-cell reference and the other as the input bulk transcriptomics data via aggregating all its single-cell gene expression profiles (Fig. S1c). Considering the cell composition of bulk RNA-seq data varies greatly in the biological circumstance, for each single-cell reference, we further changed the cell-type proportions and synthesized three corresponding bulk transcriptome data with different cell compositions. In total, 30 paired simulation data were synthesized in this study. Next, for unpaired simulations, the simulated data were derived from the above 8 scRNA-seq datasets of the human pancreas. One dataset was randomly selected as the single-cell reference data and another dataset from a different resource of the 8 scRNA-seq datasets was selected to synthesize the bulk transcriptomics data (Fig. S1d). In total, 12 unpaired simulation data were synthesized in this study.

##### *Paired and unpaired simulations for evaluation of the spatial mapping step of Bulk2Space*

For the spatial mapping step, scRNA-seq data were used to simulate spatially resolved transcriptomics. In detail, we randomly chose 10 cells from each scRNA-seq data and aggregated their gene expression profiles as a spot of pseudo spatial transcriptomics data. The spot with over 25000 UMI counts would be sampled down to 20000 UMI counts to better meet the biological situation. We also simulated pseudo spatial transcriptomics data with 100, 200, 500, 1000, and 5000 spot numbers to biologically reproduce data derived from different spatial barcoding technologies. Similar to the deconvolution step, 50 paired and 60 unpaired simulations were constructed to benchmark the mapping step of Bulk2Space.

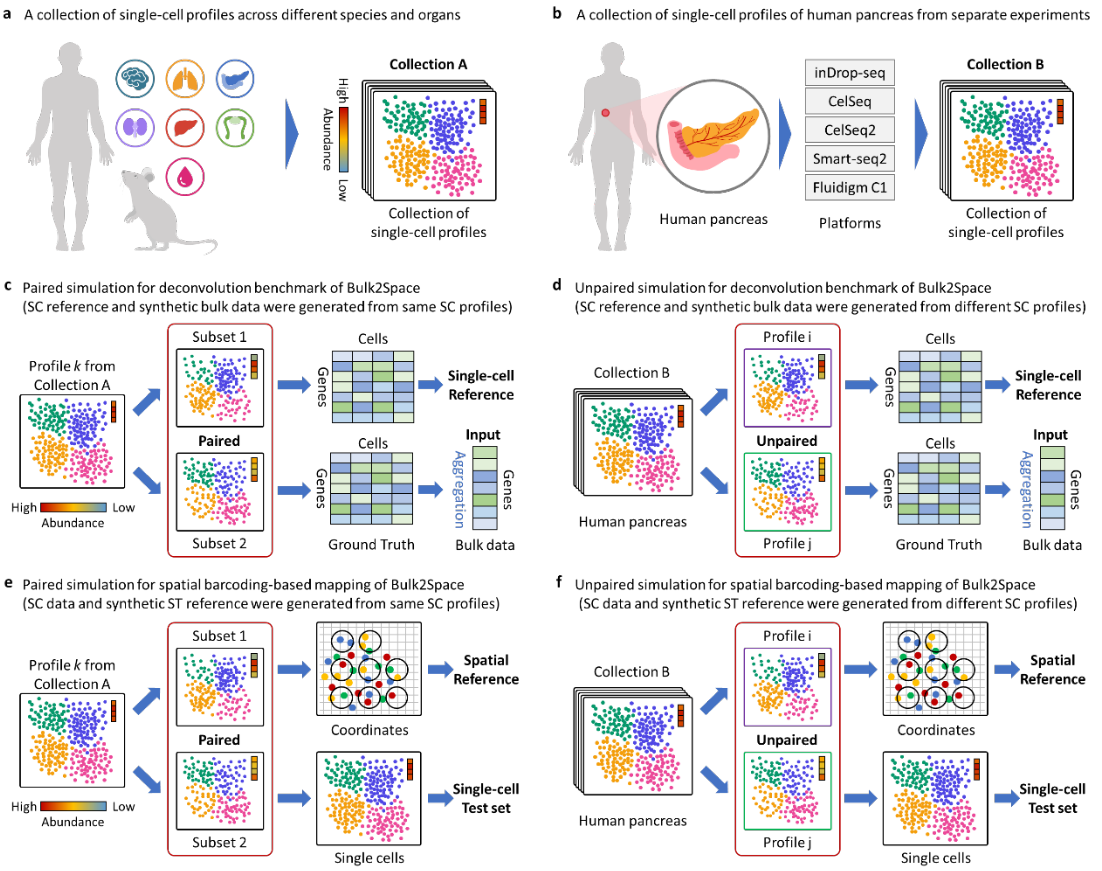

**Fig. S1. Data collection and simulation workflow**. **a**, scRNA-seq data collection for paired simulations. 10 scRNA-seq data across different species and tissues were collected. **b**, scRNA-seq data collection for unpaired simulations. 8 scRNA-seq data of the human pancreas from different platforms were collected. **c**, Paired simulation procedure for the deconvolution benchmark of Bulk2Space. Each single-cell (SC) dataset was divided into two parts, one was used as the single-cell reference and the other was aggregated to synthesize the input bulk data. **d**, Unpaired simulation procedure for the deconvolution benchmark of Bulk2Space. The reference data and the synthetic bulk data were generated from different datasets of the human pancreas. **e**, Paired simulation procedure for the spatial mapping benchmark of Bulk2Space. Each single-cell (SC) dataset was divided into two parts, one was used to simulate spatial reference via assigning single cells to pseudo spots with spatial coordinates, and the other was used as the single-cell test set. **f**, Unpaired simulation procedure for the spatial mapping benchmark of Bulk2Space. The spatial reference and the single-cell test set were generated from different datasets of the human pancreas.

#### Benchmark test for the deconvolution step of Bulk2Space

***Benchmark with paired simulations***

Thirty paired simulation data generated from 10 scRNA-seq datasets across different species and tissues were used to evaluate the performance of the deconvolution step of Bulk2Space. As shown in Fig. S2, ten out of thirty datasets were exhibited to illustrate the deconvolution performance of Bulk2Space, as supplementary information to Fig. 2a in the main text. Compared with the test set, the single-cell RNA-seq data generated by Bulk2Space showed consistent distribution (Fig. S2a). The clustering spaces of generated single-cell transcriptomics data from different datasets by Bulk2Space were exhibited in Fig. S2b.

Moreover, we compared Bulk2Space which used β-VAE as the generative model, with another two deep learning models, generative adversarial networks (GAN) and conditional generative adversarial networks (CGAN). As shown in Fig. S2c, the correlations of marker gene expression for different cell types suggested that single cells generated by Bulk2Space (beta variational autoencoder, β-VAE) and CGAN had higher correlations than GAN in gene expression between generated data and the ground truth. Although CGAN had a comparable performance as Bulk2Space in the single-cell generation, its computing speed was significantly lower than Bulk2Space. Therefore, taking time and performance into consideration, Bulk2Space chose β-VAE as the generative model for the deconvolution of bulk transcriptomics data.

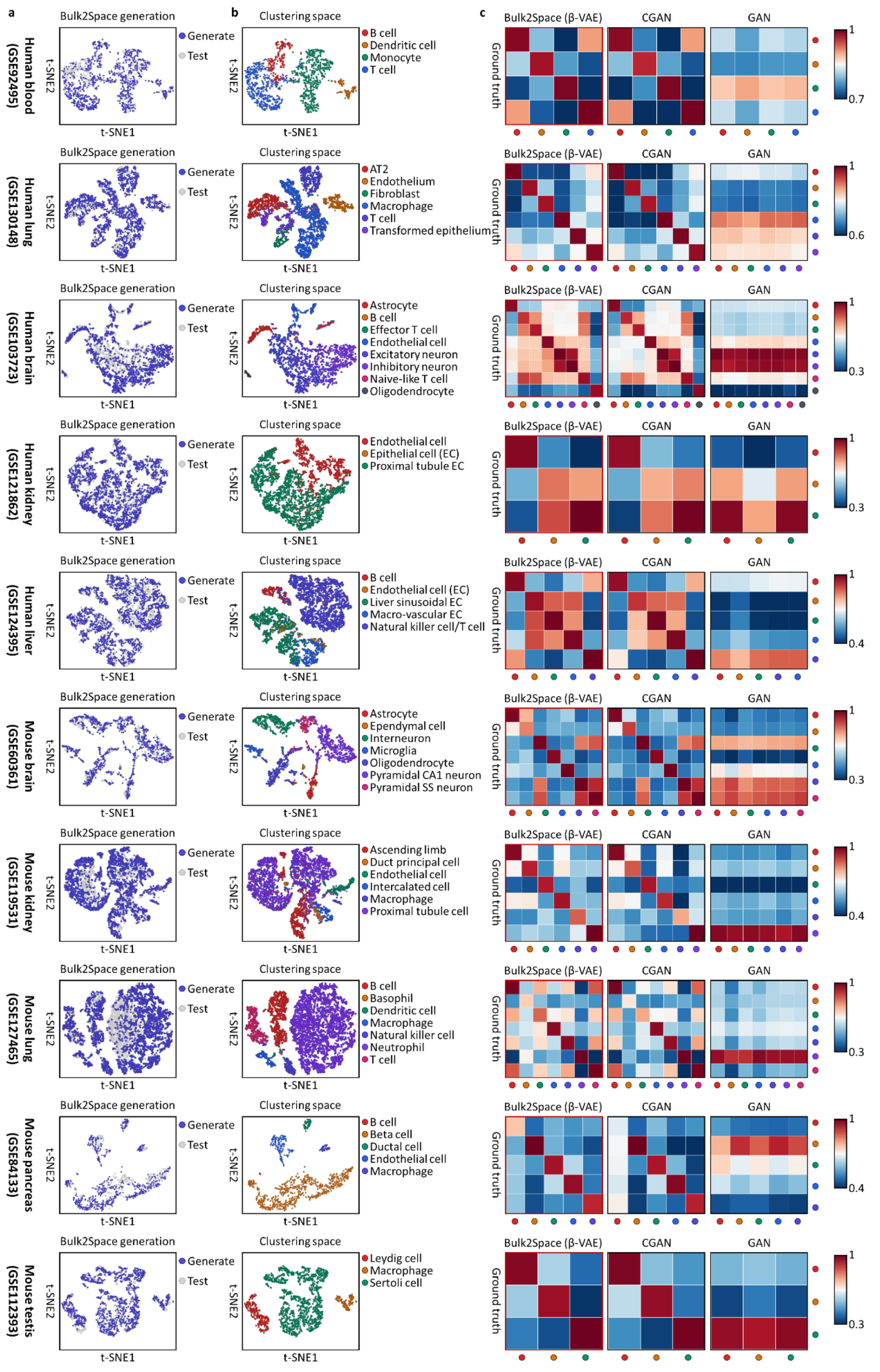

**Fig. S2**. **Deconvolution of synthetic bulk data by Bulk2Space using paired simulations. a**, The t-SNE layout showed the distribution of Bulk2Space generated single cells (blue) and the test sets (grey) in from 10 different datasets. **b**, The t-SNE plots of single cells generated by Bulk2Space demonstrated the clustering space of different cell types from 10 different datasets. **c**, Comparison of correlations of the marker gene expression between Bulk2Space (β-VAE), CGAN, and GAN. Each cell type was labeled with a color corresponded to the cell-type representation in **b**. The heatmap showed the correlation of different cell types in marker gene expression between generative methods and the ground truth. Right, the scale bar. Source data are provided as a Source Data file.

***Benchmark with unpaired simulations***

Furthermore, we benchmarked the deconvolution step of Bulk2Space using 8 separated datasets derived from the human pancreas and sequenced by different platforms. The unpaired simulation procedure was described above. In total, 12 unpaired pancreas data were simulated.

Because other deconvolution methods, such as CPM[15], CIBERSORT[16], and ImmuCC[17], can only predict cell-type proportions instead of gene expression of generated data, we compared Bulk2Space with GAN, CGAN, and a Bayesian deconvolution method termed bMIND[18]. As illustrated in Fig. S3a, the gene expression correlation between the generated single-cell data and the ground truth of the constructed bulk data was calculated to evaluate and compare the four candidate algorithms. Bulk2Space outperformed the other three methods by the *Pearson* correlation of gene expression and the gene expression variation (root mean squared error, RMSE). We found that bMIND showed a higher *Spearman* correlation of gene expression than Bulk2Space, GAN, and CGAN. Then we compared the correlations of marker gene expression for different cell types between four generations and the ground truth (Fig. S3b). The results suggested that single cells generated by Bulk2Space and CGAN had higher correlations than GAN and bMIND in gene expression between generated data and the ground truth.

Since the unpaired simulation data were derived from different platforms. We discuss whether the quality, for instance, sequencing depth, of the reference data could affect the deconvolution of Bulk2Space. Compared with paired simulation, the overall correlation of gene expression between generated data and ground truth decreased because of batch effects. However, when using unpaired simulations, as shown in Fig. S3, the comparison between 12 simulations remained robust, suggesting that Bulk2Space was not so dependent on the single-cell reference, although the quality of the single-cell reference directly affected the characterization of the clustering space of different cell types.

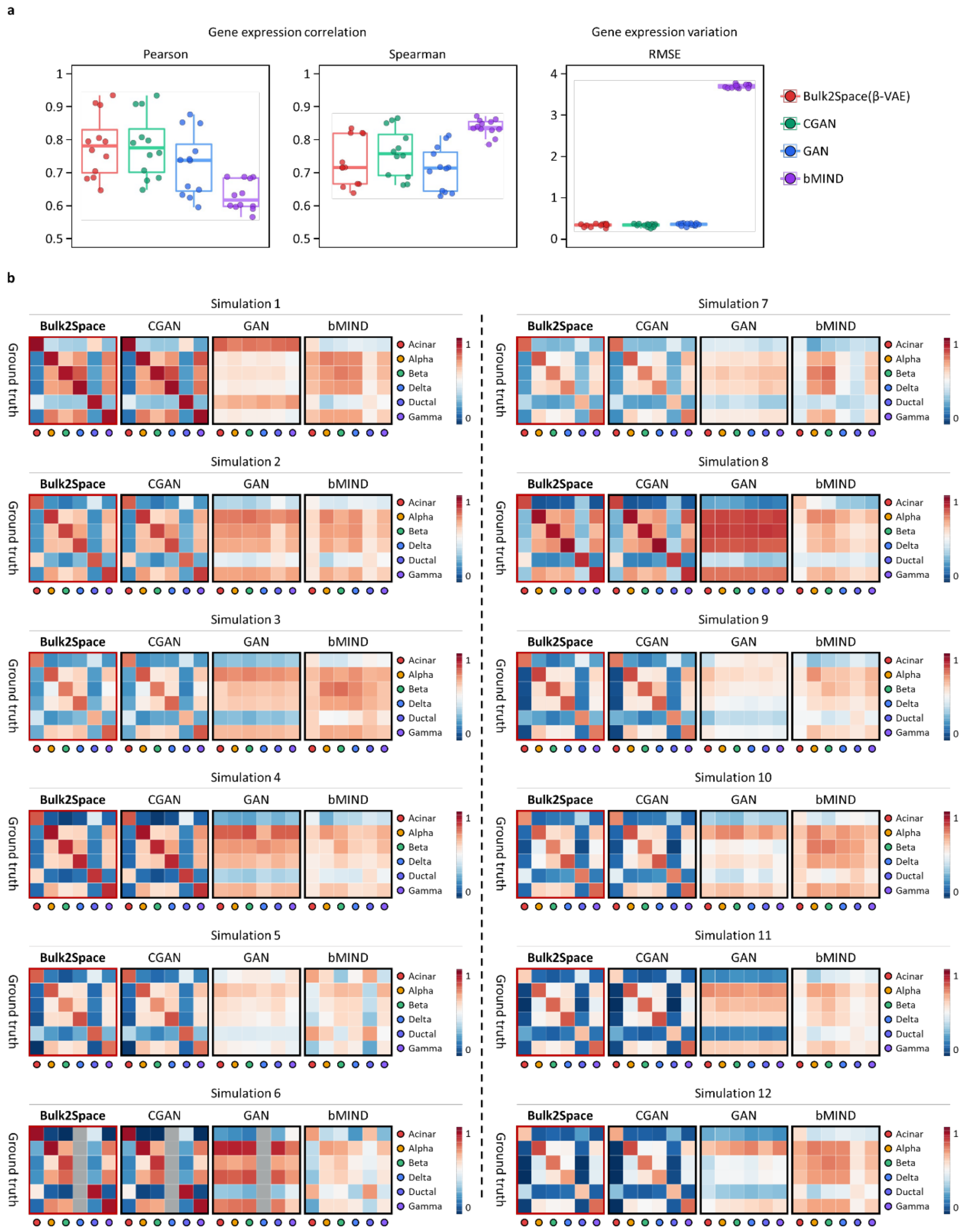

**Fig. S3**. **Deconvolution of synthetic bulk data by Bulk2Space using unpaired simulations and comparison with other methods. a**, Comparison of the performance of deconvolution between four different methods using unpaired simulation data (n=12). Left, *Pearson* correlation of gene expression between generated and input bulk data. Middle, *Spearman* correlation of gene expression between generated and input bulk data. Right, gene expression variation between generated and input bulk data using RMSE. Data are presented as boxplots (minima, 25th percentile, median, 75th percentile, and maxima). **b**, Comparison of correlations of the marker gene expression between Bulk2Space (β-VAE), CGAN, GAN, and bMIND. The heatmap showed the correlation of different cell types in marker gene expression between generative methods and the ground truth. Right, the scale bar. Source data are provided as a Source Data file.

##### ***Noise introduction for robustness evaluation of Bulk2Space***

To further investigate the robustness of Bulk2Space, we introduced two noise mechanisms to test the performance of the algorithm. One mechanism changes the expression values of certain genes in randomly selected cells (Fig. S4a), and the other alters the cell type labels of selected cells (Fig. S4b). In both cases, the correlations in gene expression and cell types between the generated and test data decreased with the increasing noise, while the RMSEs gradually increased. As illustrated in Fig. S4c and Fig. S4d, for the first type of noise introduction, the gene expression correlation decreased dramatically even when the noise was at a lower extent, while the cell type correlation remained relatively stable. This is because cell types are usually determined by a group of marker genes, and the variation of individual genes has a relatively low impact on the determination of cell identity. However, the variation of genes can cause huge changes in the value of gene expression when the single-cell profiles are generated. The results demonstrated that Bulk2space could effectively avoid the overfitting phenomenon. In contrast, for the second type of noise introduction, when cell types were mislabeled, the gene expression correlation and RMSE remained steady, yet the cell type correlation decreased rapidly when the noise increased. As expected, altering the cell type labels will not change the cell gene expression, but it can affect the characterization of the clustering space. Therefore, when generating a single-cell expression profile, the resulting single cell may locate between different clustering spaces, thus causing the cell to be assigned to the wrong cell type. However, the overall expression level of genes remained relatively stable.

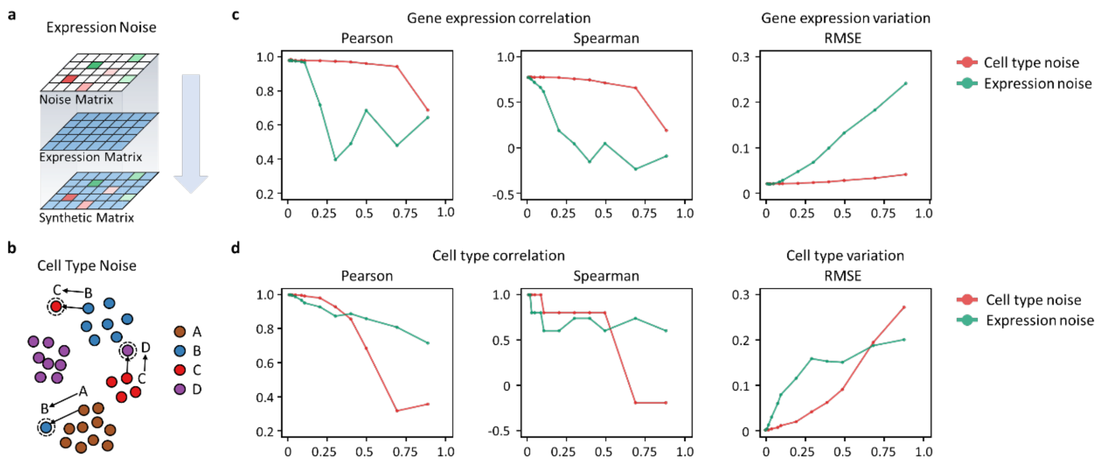

**Fig. S4**. **Noise introduction for the robustness evaluation of Bulk2Space**. **a**, Synthesis of expression noise data. A noise matrix was composed of zero values and a certain number of variation values, red donated up-regulation of gene, and green represented down-regulation. The noise matrix was added to the original expression matrix to synthesize the single-cell gene expression profiles with expression noise introduction. **b**, Synthesis of cell type noise data. A certain number of single cells were randomly selected and their labels were replaced with other cell types to generate new single-cell profiles with cell type noise introduction. **c** and **d**, Gene expression correlation and RMSE, and cell type correlation and RMSE for noise data and test set. Left, Pearson correlation, middle, Spearman correlation, right, RMSE. Expression noise is colored in green and cell type noise is colored in red. Source data are provided as a Source Data file.

***Deconvolution of bulk RNA-seq data using annotation-free single-cell reference by Bulk2Space***

In this section, we performed the deconvolution step of Bulk2Space using single-cell reference[1] without cell-type annotations. As shown in Fig. S5, we first clustered the reference single-cell profiles into five clusters using the existing clustering algorithm Louvain of Seurat[19]. Based on the clustering space of the five resulting clusters, single cells were generated from simulated bulk data by Bulk2Space. Compared with the test set, the single-cell RNA-seq data generated by Bulk2Space showed consistent distribution (Fig. S5a). The clustering space of generated single-cell transcriptomics data and the reference data were exhibited in Fig. S5b. And the correlation of marker gene expression for different cell clusters showed that the corresponding cell clusters had a higher correlation, suggesting that Bulk2Space can robustly generate single cells using unannotated single-cell reference (Fig. S5c).

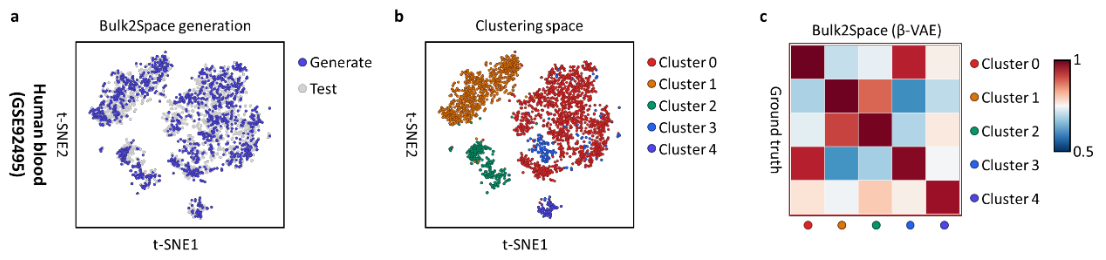

**Fig. S5**. **Deconvolution of bulk data using unannotated single-cell reference by Bulk2Space**. **a**, The t-SNE layout showed the distribution of Bulk2Space generated single cells (blue) and the test sets (grey). **b**, The t-SNE plots of single cells generated by Bulk2Space demonstrated the clustering space of different cell clusters. **c**, Correlation of the marker gene expression for different clusters between Bulk2Space generated single-cell data and the ground truth. Right, the scale bar. Source data are provided as a Source Data file.

***Perturbation analysis of Bulk2Space***

Bulk2Space utilized β-VAE for single-cell generation. β-VAE is a modification of the Variational Autoencoder with a special emphasis on discovering disentangled latent factors. For example, a model trained on photos of human faces might capture the gentle skin color, hair length, hair color, emotion, and many other relatively independent factors in separate dimensions. One benefit that often comes with disentangled representation is good interpretability and straightforward generalization to various tasks. Intuitively, this latent space makes beta-VAE an effective tool for generating and understanding variations in natural data. We conducted perturbation analysis with β-VAE. Since the original latent vectors obey the Gaussian distribution, here we changed the distribution to uniform distribution and Poisson distribution. As illustrated in the Fig. S6, cells generated under the original setting (Gaussian distribution) were mapped closer to the original cells (Fig. S6a), while under the uniform (Fig. S6b) and Poisson (Fig. S6c) distributions, the generated cells are more clustered, resulting in poor clustering performance. This confirmed that better generation results can be obtained by assuming that the latent vectors follow the Gaussian distribution. The dimensionality of the latent space learned by the beta-VAE is 256.

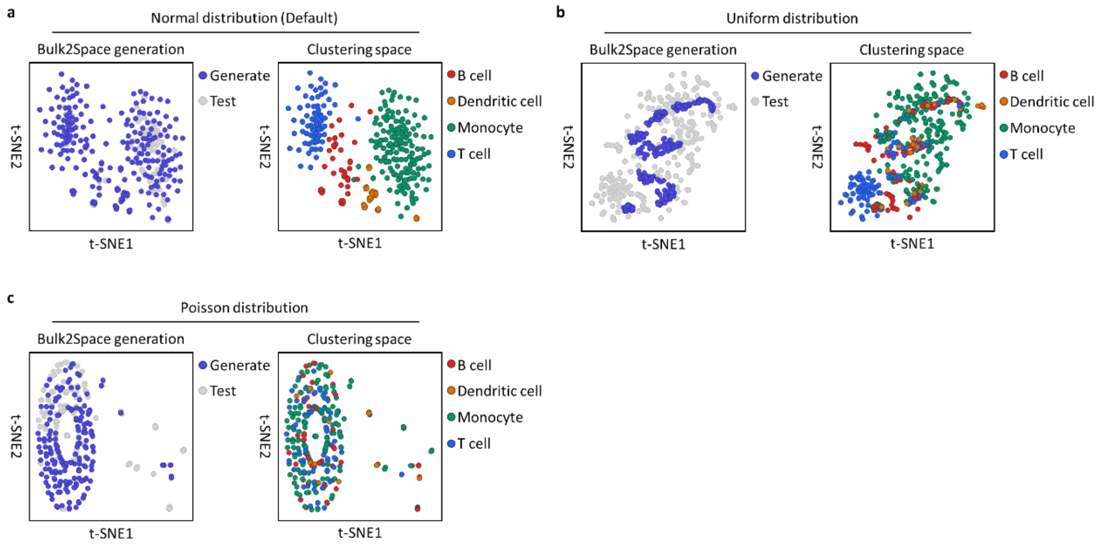

**Fig. S6**. **Perturbation analysis of Bulk2Space**. **a**, Single-cell generation using Gaussian distribution. Left, distribution of Bulk2Space generated single cells (blue) and the test set (grey). Right, clustering space of the generated single cells. **b**, Single-cell generation using Uniform distribution. **c**, Single-cell generation using Poisson distribution.

##### *Deconvolution performance evaluation of Bulk2Space using paired biological bulk and single-cell datasets*

Based on the robust performance of Bulk2Space on simulated data, we further validated it with biological data. In biological circumstances, bulk and single-cell RNA-seq data from the same tissue can be treated as paired datasets because they share identical conditions. In this study, three paired bulk and single-cell RNA-seq data of different mice liver (GSE119340)[20] fed with standard chow were collected to evaluate the performance of Bulk2Space. As shown in Fig. S7, three paired mice bulk and single-cell profiles were used as input bulk data and single-cell reference data. Through Bulk2Space deconvolution, the correlation of the marker gene expression for different cell types between generated and the reference single-cell profiles showed that Bulk2Space could be well applied in biological scenarios.

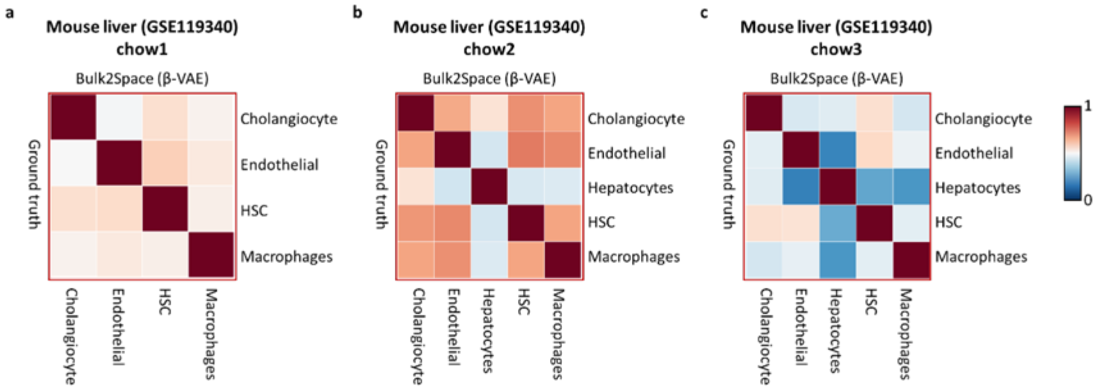

**Fig. S7**. **Deconvolution of bulk data using biological datasets by Bulk2Space**. **a**, **b**, and **c**, Correlation of the marker gene expression for different clusters between Bulk2Space generated single-cell data and the reference data. Right, the scale bar. Source data are provided as a Source Data file.

#### Benchmark test for the spatial mapping step of Bulk2Space

##### *Benchmark with paired simulations for spatial barcoding-based reference*

As described above, paired simulation data were used to evaluate the performance of the spatial mapping step of Bulk2Space for spatial barcoding-based reference, as the supplementary information for Fig. 2c in the main text. Each scRNA-seq data was randomly separated into two parts, with one as the single-cell data and the other as the spatial reference via assigning 10 cells to each spot and aggregating their expression. Cells in the single-cell data were mapped to spots based on the pairwise similarity of gene expression. Therefore, Bulk2Space could provide spatially resolved single-cell transcriptomics data. The comparison of the cell-type composition for each spot between Bulk2Space and the spatial reference across 10 example datasets was shown in Fig. S8. The spatial composition of cell types predicted by Bulk2space was very close to the spatial reference.

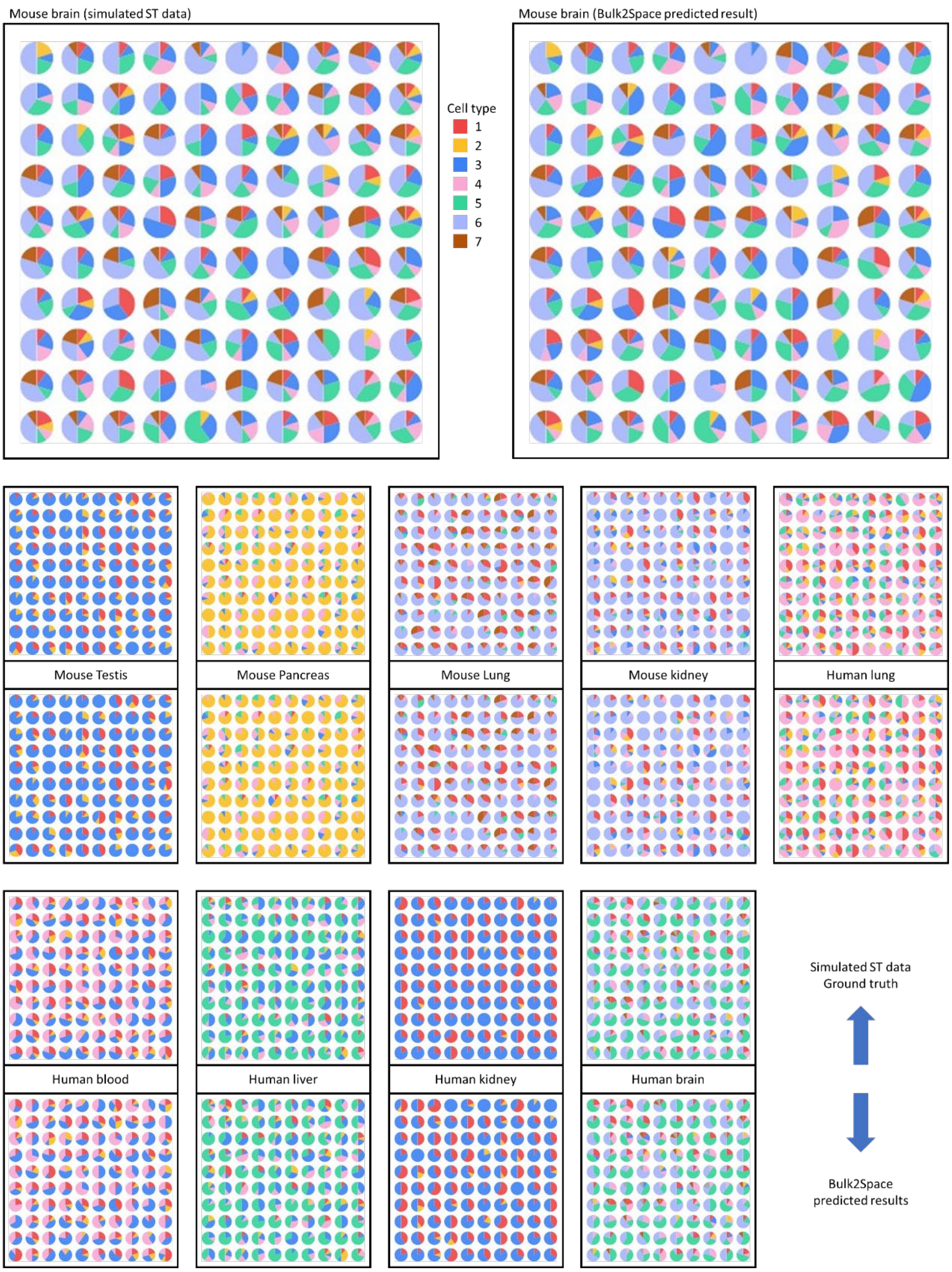

**Fig. S8**. **Comparison of the cell-type composition of each simulated spot between Bulk2Space and the spatial reference using paired simulations**. First row, Left, cell-type spatial composition of simulated spatial reference data of the mouse brain. Right, spatial composition of cell types of the mouse brain predicted by Bulk2Space. Others, Top, cell-type spatial composition of simulated spatial reference data of different tissues. Bottom, spatial composition of cell types of different tissues predicted by Bulk2Space. Source data are provided as a Source Data file.

***Benchmark with unpaired simulations for spatial barcoding-based reference***

Also, we used unpaired simulations to evaluated the the spatial mapping of Bulk2Space as the supplementary information for Fig. 2d in the main text. The single-cell and spatial reference data were selected from separated datasets of the human pancreas. The comparison of the cell-type composition for each spot between Bulk2Space and the spatial reference was shown in Fig. S9.

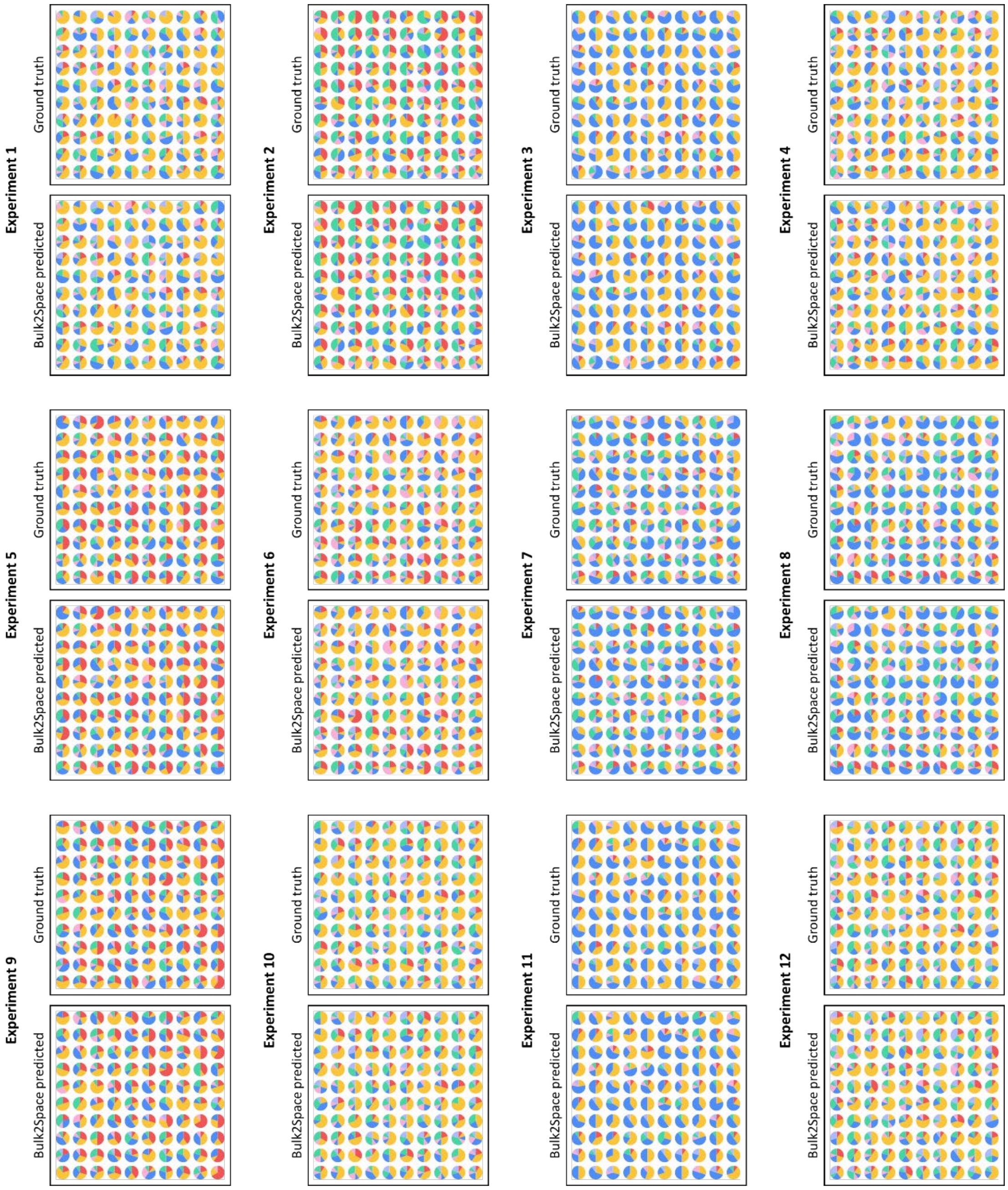

**Fig. S9**. **Comparison of the cell-type composition of each simulated spot between Bulk2Space and the spatial reference using unpaired simulations**. Top, cell-type spatial composition of simulated spatial reference data. Bottom, spatial composition of cell types predicted by Bulk2Space. Source data are provided as a Source Data file.

***Leave-one-out test for the spatial mapping step of Bulk2Space with image-based methods***

The MERFISH dataset containing 150 target genes was used as the spatial reference. Two sets of single-cell data, one from the same experiment as the MERFISH and the other from a separated experiment, were used as paired and unpaired single-cell data, respectively. Since MERFISH and single-cell data were heterogeneous, a joint analysis of the two kinds of data was carried out in advance. Finally, a leave-one-out test was carried out.

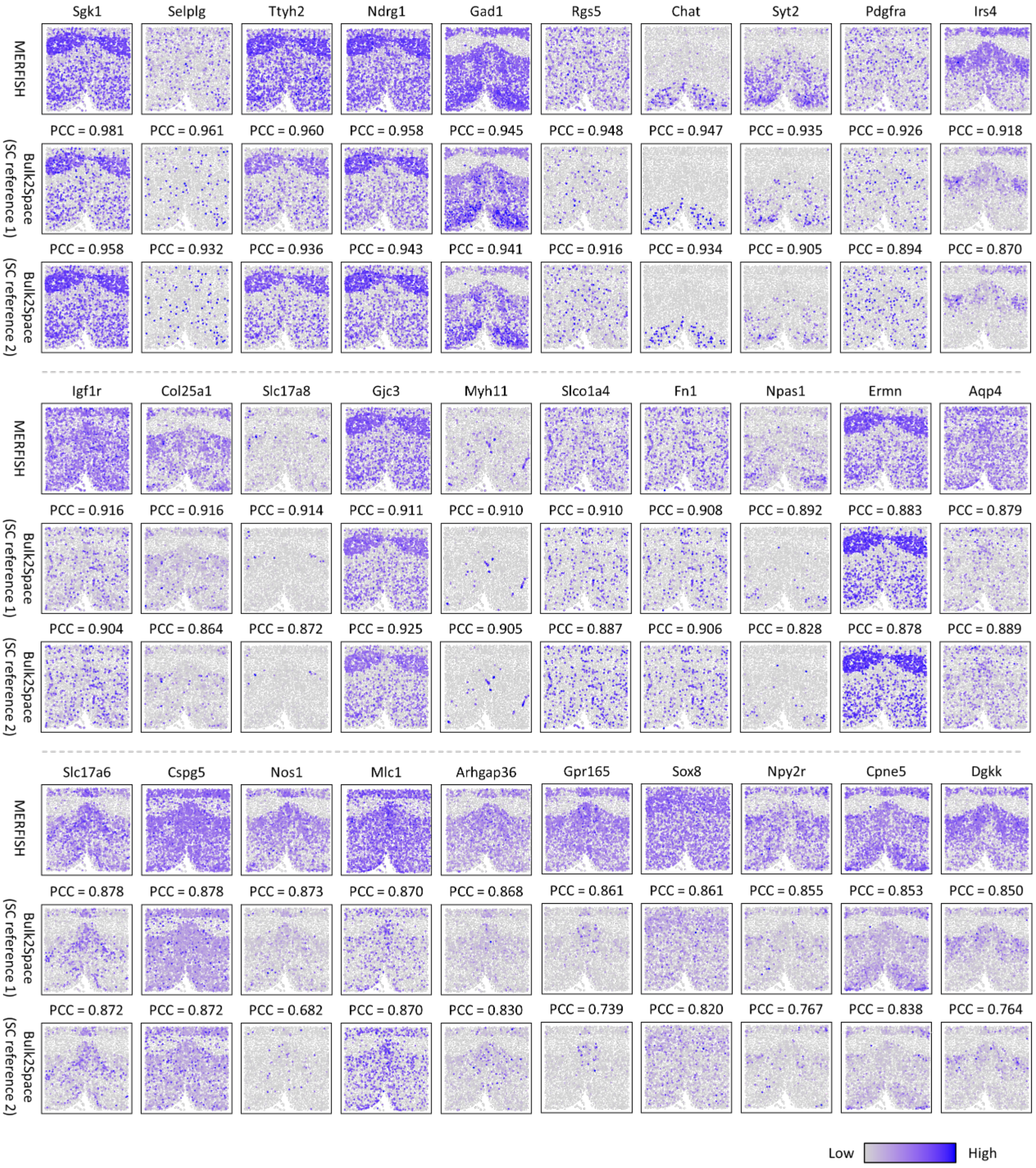

**Fig. S10**. **Comparison of the spatial expression of 30 genes between MERFISH and Bulk2Space results with the paired and unpaired single-cell data using leave-one-out test**. Top, spatial expression of genes in MERFISH. Middle, spatial expression of genes predicted by Bulk2Space using paired single-cell data. Bottom, spatial expression of genes predicted by Bulk2Space using unpaired single-cell data. *Pearson* correlation coefficients (PCC) of the spatial gene expressions are listed.

Source data are provided as a Source Data file.

In detail, 149 genes in MERFISH were utilized to map single cells to coordinates based on the pairwise similarity between single-cell and spatial reference profiles, and the remaining 1 gene was used for the validation of predicted spatial expression. As shown in Fig. S10, the spatial expressions of 30 genes predicted by Bulk2Space for the paired and unpaired single-cell data were compared with MERFISH data. The spatial mapping result of paired single-cell data was better than that of unpaired single-cell data in terms of the PCC score for each gene.

***Five-fold cross-validation for the spatial mapping step of Bulk2Space with image-based methods***

We further verified Bulk2Space's spatial mapping using five-fold cross-validation for image-based methods (Fig. S11).

Specifically, the 150 targeted genes in MERFISH were split into five folds using R package caret (version 6.0-92), and 80% of genes were used as the referece and the remaining 20% were used for validation of predicted spatial expression. As shown in Fig. S11, the spatial expressions of 25 genes predicted by Bulk2Space for the paired and unpaired single-cell data were compared with MERFISH data.

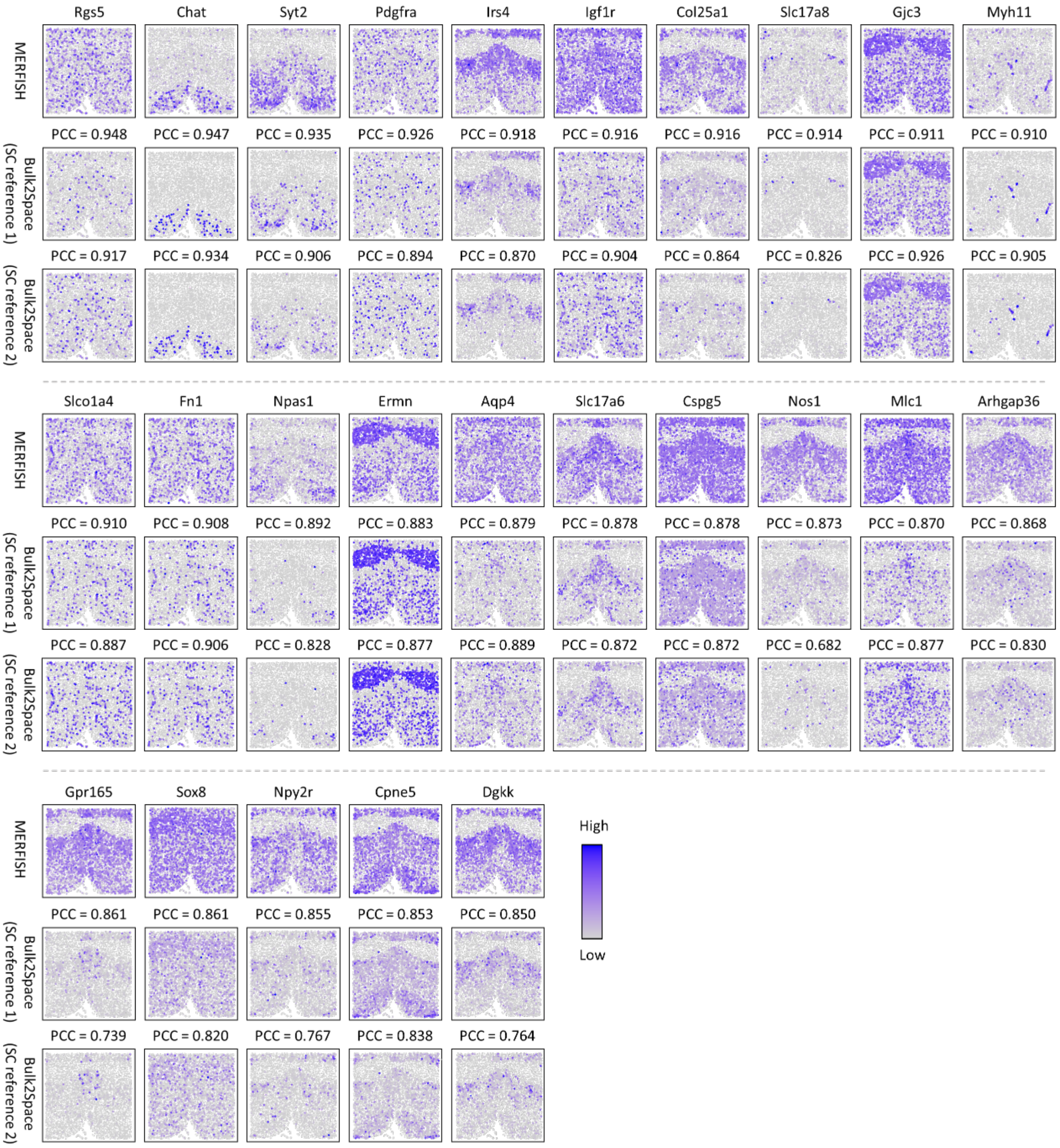

**Fig. S11**. **Comparison of the spatial expression of 25 genes between MERFISH and Bulk2Space results with the paired and unpaired single-cell data using an 80%~20% cross-validation**. Top, spatial expression of genes in MERFISH. Middle, spatial expression of genes predicted by Bulk2Space using paired single-cell data. Bottom, spatial expression of genes predicted by Bulk2Space using unpaired single-cell data. PCC of the spatial gene expressions were listed. Source data are provided as a Source Data file.

##### *Performance evaluation of the spatially mapping step of Bulk2Space using biological datasets*

To validate the spatial mapping step of Bulk2Space in the real situation, two biological datasets, a spatial reference, and a single-cell data, from separated experiments were used to allocate single cells onto the spatial coordinates of the mouse hippocampus. The spatial reference data were sequenced by Slide-seq v2[21] and the scRNA-seq data[22] were downloaded from the Seurat website (https://satijalab.org/seurat/articles/spatial_vignette.html). We down-sampled the Slide-seq v2 data to 5000 spots to speed up the spatial mapping step.

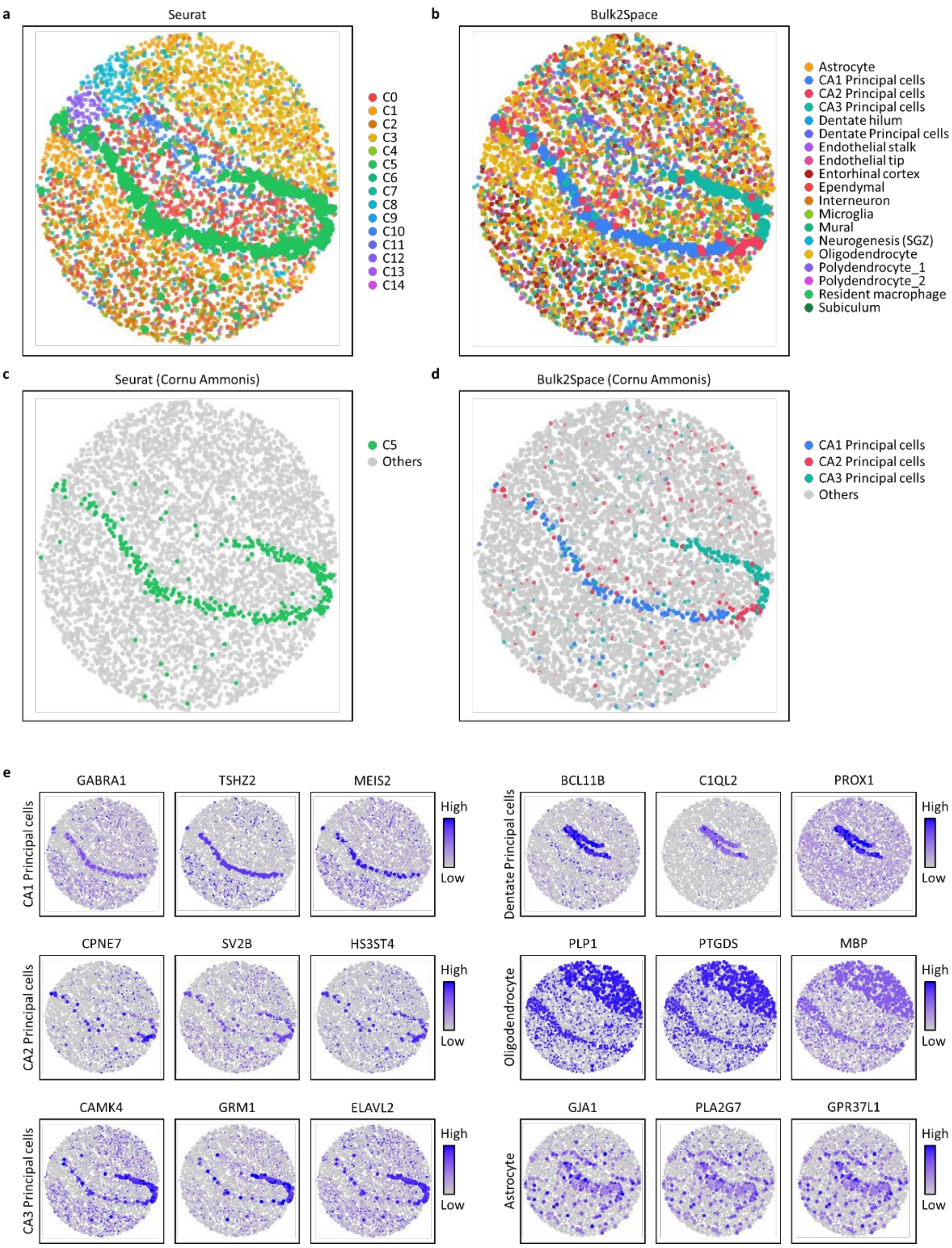

**Fig. S12**. **Reconstruction of the mouse hippocampus structure at single-cell resolution using biological datasets**. **a**, Spatial clustering of the spots by Seurat. **b**, Spatial mapping of individual cells in the scRNA-seq data to spots in the spatial reference by Bulk2Space. **c**, Annotations of spots in CA1, CA2, and CA3 subregions by Seurat. **d**, Annotations of single cells that were mapped to spots in CA1, CA2, and CA3 subregions by Bulk2Space. **e**, Spatial expression distribution of three marker genes of CA1 principal cells, CA2 principal cells, CA3 principal cells, dentate principal cells, oligodendrocytes, and astrocytes. Source data are provided as a Source Data file.

As shown in Fig. S12, for comparison, we first performed the Seurat clustering algorithm on the spatial data (Fig. S12a). Notably, Seurat only provided a classified the spots directly into 15 clusters based on the expression profiles without annotating their cell types. Then, we performed Bulk2Space's spatial mapping step to calculate the pairwise similarity of cells and spots between the scRNA-seq and the spatial reference data. Each spot was deconvolved into single cells with cell-type annotations (Fig. S12b). The spatial distribution of single cells predicted by Bulk2Space was consistent with the real structural pattern of cell types in the mouse hippocampus region.

Subsequently, we highlighted the CA1, CA2, and CA3 subregions in the mouse hippocampus. Seurat annotated the three subregions with the same labels (Fig. S12c). In contrast, Bulk2Space successfully reconstructed the refined structure of the three subregions by mapping the corresponding cell types to these regions (Fig. S12d).

Moreover, we investigated the spatial expression patterns of marker genes for six spatial specific cell types, CA1 principal cells, CA2 principal cells, CA3 principal cells, dentate principal cells, oligodendrocytes, and astrocytes. As shown in Fig. S12e, the expression patterns of the marker genes showed obvious spatial distribution characteristics, which were consistent with the spatial distributions of the corresponding cell types. The results suggested that Bulk2Space could map cells to spatial locations precisely in biological situations.

#### Robustness evaluation of Bulk2Space using repeated data

As described in the main text, Bulk2Space employed β-VAE to generate single-cell profiles within the clustering space of cell types. Here, we discuss whether the randomness of β-VAE could affect the generation of single-cell data.

***Robustness evaluation of the deconvolution step using 100 repetitions of single-cell generation***

Here, we conducted 100 repetitions on the deconvolution step of Bulk2Space to evaluate the robustness of β-VAE using the human peripheral blood[1] scRNA-seq data.

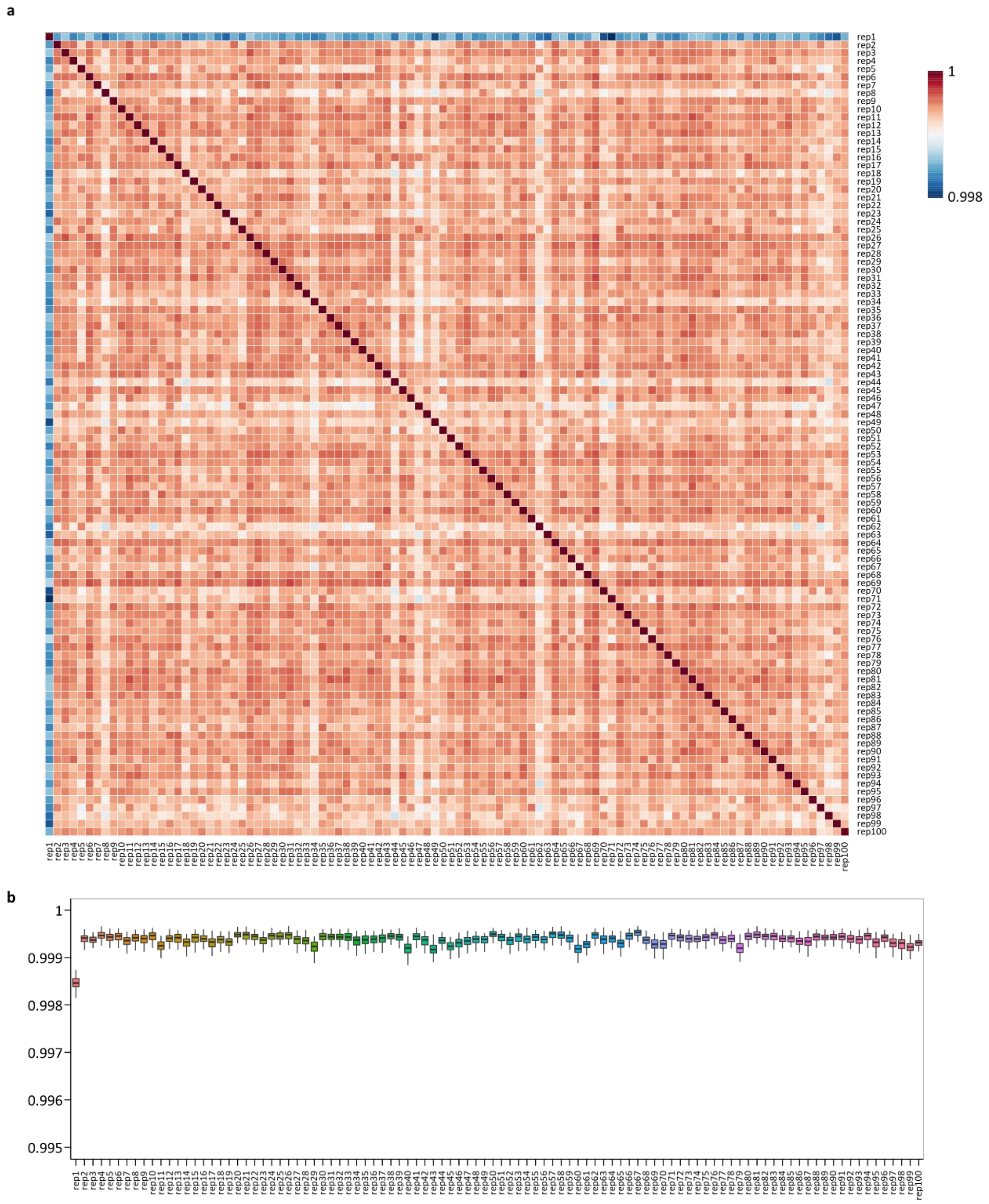

**Fig. S13**. **Robustness evaluation of the deconvolution step of Bulk2Space with 100 repetitions of β-VAE**. **a**, Pairwise correlation of generated single-cell profiles between different replicates of deconvolution. **b**, Averaged correlation of generated single-cell profiles between each replicate and other replicates. The number of replicates is 99. Data are presented as boxplots (minima, 25th percentile, median, 75th percentile, and maxima). Source data are provided as a Source Data file.

The data simulation procedure was consistent with the earlier statement. As shown in Fig. S13, the single-cell generation remained highly robust across 100 repetitions of β-VAE, with the pairwise correlations of generated single-cell profiles between replicates above 0.998 (Fig. S13a). For each replicate, the averaged correlation of the generated single-cell data with other replicates remained consistent.

***Robustness evaluation of the spatial mapping step on two spatial barcoding-based references using three repetitions of single-cell generation***

With the robust performance on the single-cell generation in the deconvolution step, we further investigated whether the randomness of the single-cell generation would affect the spatial mapping results.

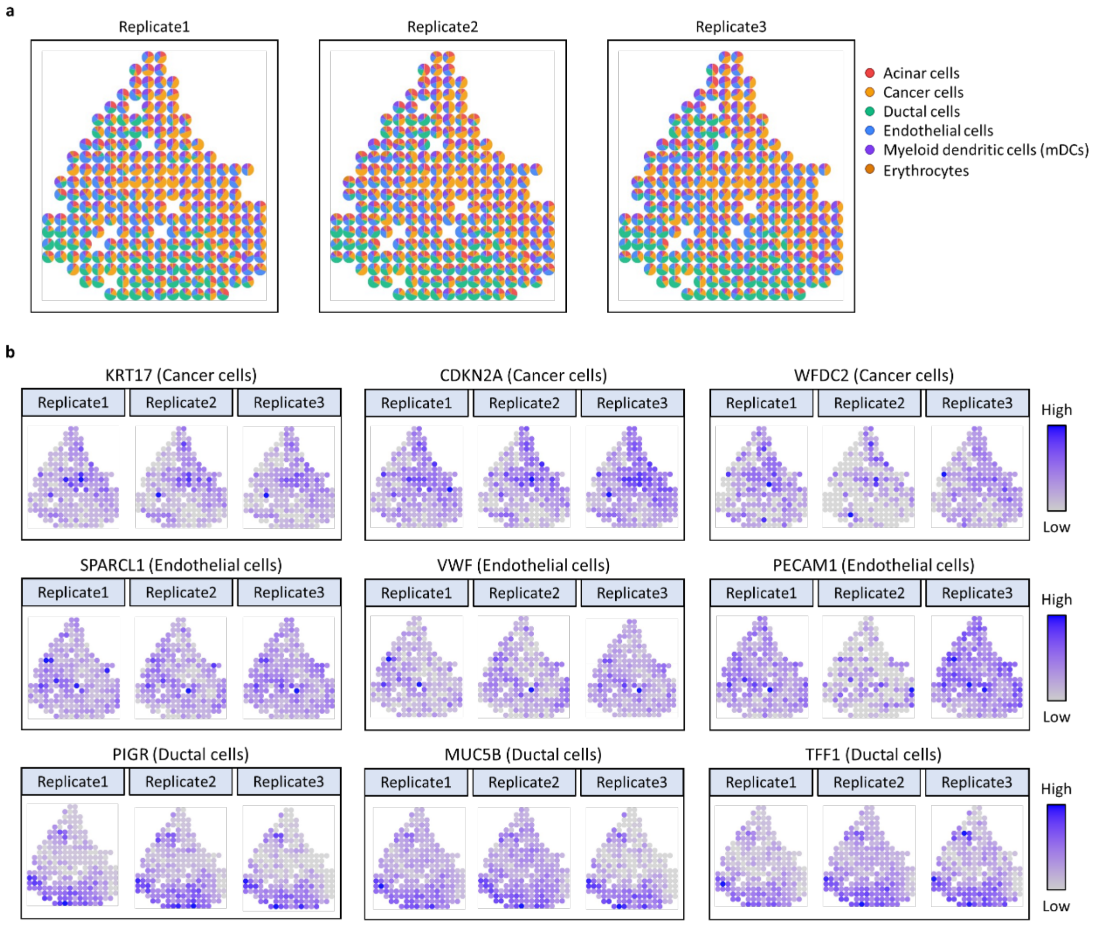

**Fig. S14**. **Spatial deconvolution of pancreatic ductal adenocarcinoma (PDAC) data by Bulk2Space using 3 repetitions**. **a**, Spatial deconvolution of the three replicates of bulk PDAC data. Each spot represented the composition and proportion of cell types. **b**, Spatial expression patterns of cell-type-specific marker genes for cancer cells, endothelial cells, and ductal cells. Source data are provided as a Source Data file.

We mapped single cells generated from three replicates of the deconvolution of simulated bulk PDAC data[23] to spatial coordinates by Bulk2Space. As shown in Fig. S14a, the spatial distribution and composition of cell types in the three repeated experiments were generally very similar. Moreover, the spatial expression of cell type-specific genes matched the spatial distribution of corresponding cells (Fig. S14b).

Similarly, we performed Bulk2Space to map single cells generated from three replicates of the deconvolution of simulated bulk melanoma data[24, 25] to spatial coordinates. As shown in Fig. S15, the spatial deconvolution results suggested a robust performance of Bulk2Space with 3 repetitions.

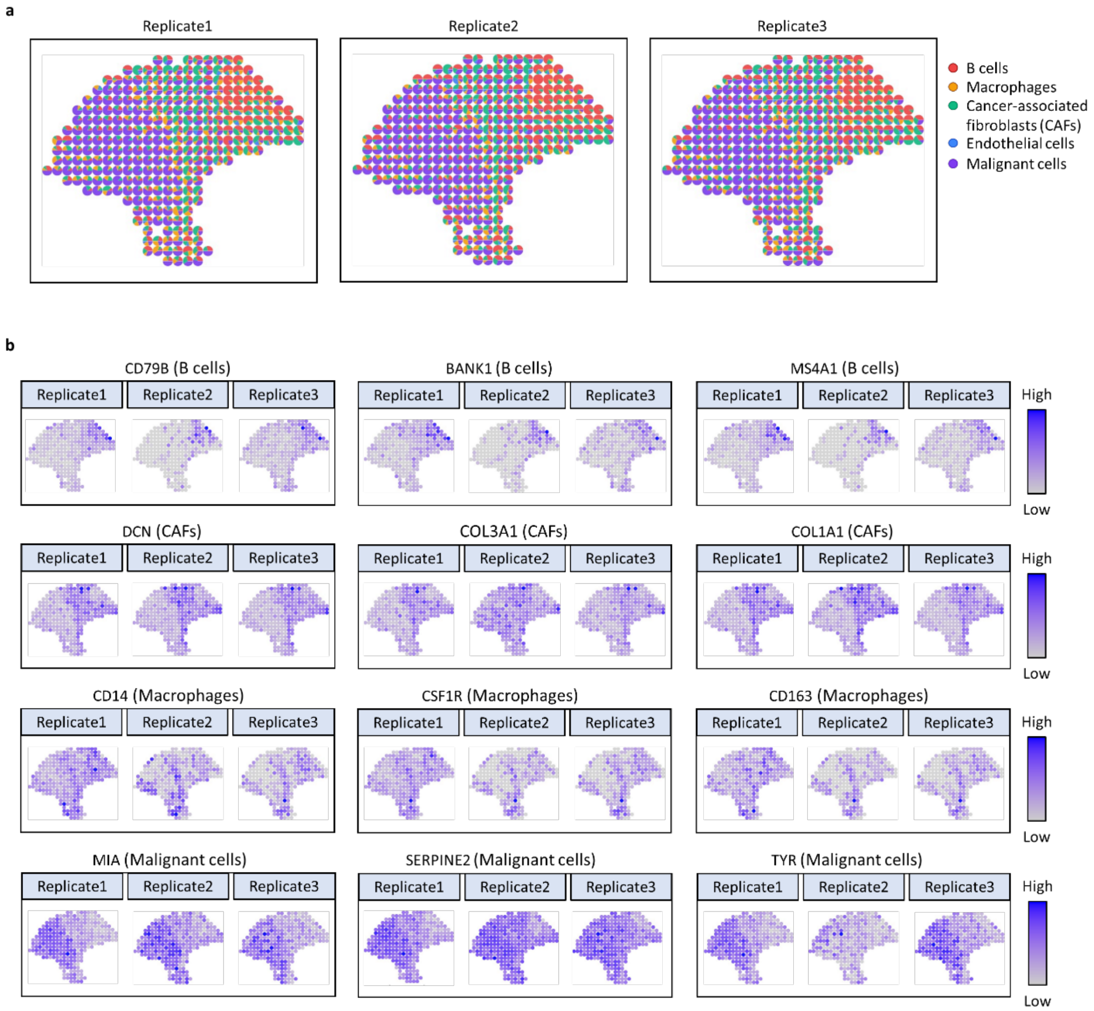

**Fig. S15**. **Spatial deconvolution of melanoma data by Bulk2Space using 3 repetitions**. **a**, Spatial deconvolution of the three replicates of bulk melanoma data. Each spot represented the composition and proportion of cell types. **b**, Spatial expression patterns of cell-type-specific marker genes for B cells, cancer-associated fibroblasts (CAFs), and malignant cells. Source data are provided as a Source Data file.

In conclusion, the repetition test suggested that the randomness of the single-cell generation function in Bulk2Space would affect the deconvolution and spatial mapping results to a small extent. However, although the single-cell data generated each time were slightly different, the overall prediction results showed robust performance of Bulk2Space in the spatial distribution of cell types, the cell-type composition and proportion in spots, and the spatial patterns of gene expression.

### Validation of Bulk2Space with biological data

#### Bulk2Space reveals spatial, molecular, and functional heterogeneity of B cells in melanoma using consecutive slices

##### *Spatial analysis of bulk melanoma by Bulk2Space*

To further verify the performance of Bulk2Space, two consecutive slices were used to demonstrate the spatial deconvolution result. The melanoma data were derived from Tirosh[24] et al. and Thrane[25] et al., a single-cell transcriptomics data and a spatially resolved transcriptomics data of the melanoma, respectively. The scRNA-seq data were used as a single-cell reference. For spatial transcriptomics data, one slice of tissue sections was used as the spatial reference (Slice 1), and another slice from a separated tissue (Slice 2) was used for bulk data synthesis (Fig. S16a).

The expression correlation of marker genes for five cell types between Bulk2Space result and Slice 2 was shown in Fig. S16b. After deconvolution, generated single cells were mapped to the coordinates of spots in Slice 1 by Bulk2Space (Fig. S16c). As illustrated in Fig. S16d, the composition of cell types in each spot of the tissue showed a strong spatial pattern, which was consistent with the histological annotation of the tissue section. The spatial distribution of each cell type is shown in Fig. S16e. The predicted results for B cells, cancer-associated fibroblasts (CAFs), macrophages, and malignant cells showed distinct spatial heterogeneity. Next, we compared the expression of marker genes in both predicted and reference tissues (Fig. S16f) and found that the spatial pattern of the marker genes in the predicted tissue was highly correlated with that in the spatial reference (Fig. S16g).

##### *Bulk2Space revealed the spatial heterogeneity of B cells*

Surprisingly, B cells assigned to the melanoma area were quite different from those assigned to the lymph node area (Fig. S16h), which suggested that there was spatial heterogeneity in B cells from different tissue regions. However, this spatial difference cannot be detected by other deconvolution methods that only predict the composition of cell types in each spot. The differential expression analysis (Fig. S16i) and pathway enrichment (Fig. S16j) were carried out to further investigate the molecular and functional heterogeneity of B cells. As shown in Fig. S16j, B cells in the lymph node area were related to immunoregulation such as lymphocyte activation, T cell activation, and immune effector process, etc. However, in the melanoma region, many pathways, such as regulation of growth, cell-substrate junction, cell-substrate adhesion, and other pathways associated with cancer cell growth, adhesion, and diffusion were enriched**.** The spatial variation of B cells illustrated the molecular architecture of the melanoma microenvironment.

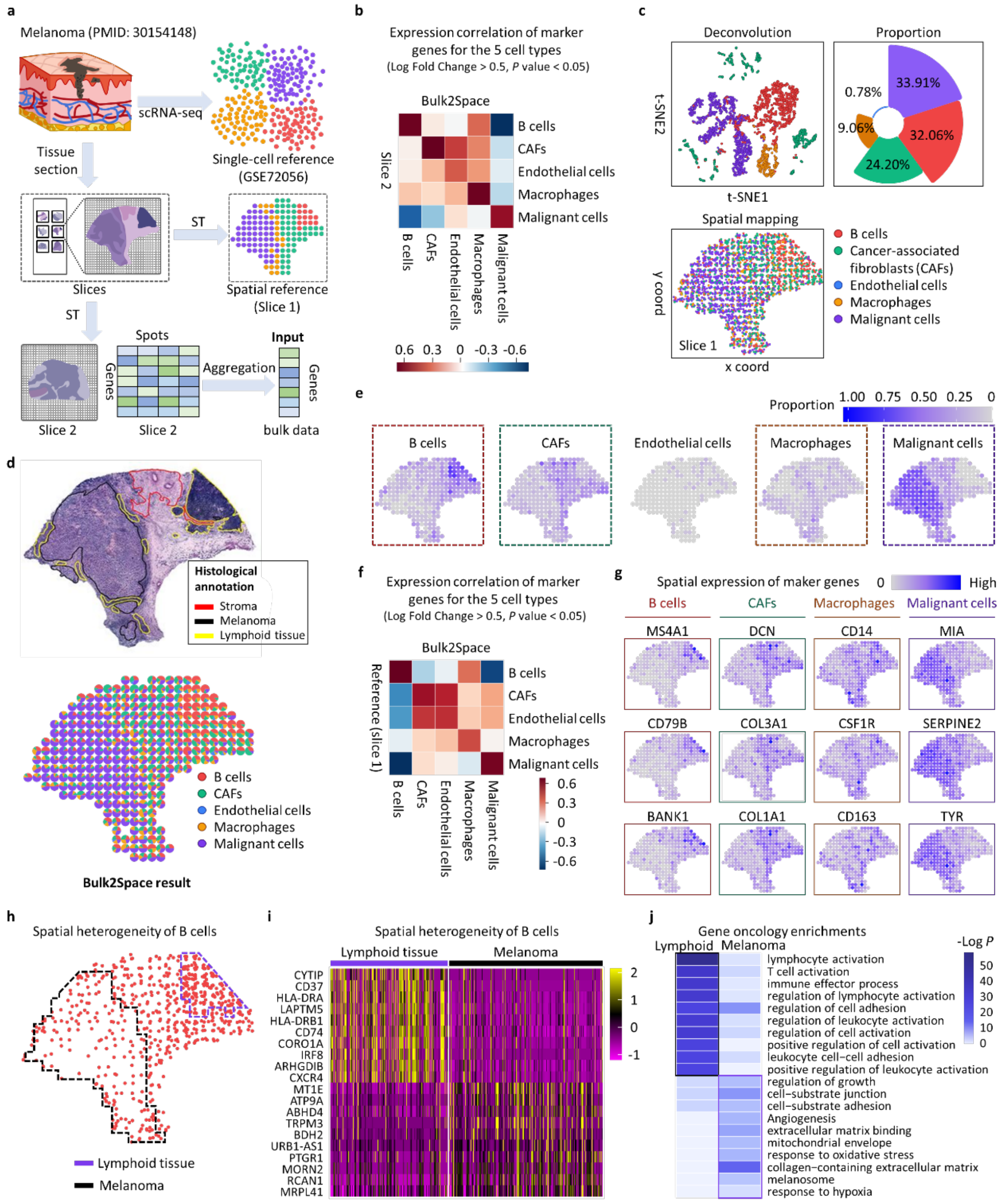

**Fig. S16. Spatial deconvolution of the melanoma by Bulk2Space using discrete slices. a**, Synthesis of the bulk transcriptome data. The scRNA-seq data is used as the deconvolution reference, one slice of the sectioned tissue is employed as the spatial reference (Slice 1), and another slice derived from a separated tissue (Slice 2) is used to synthesize the input bulk transcriptomics data. **b**, Expression correlation of marker genes for five predicted cell types between Bulk2Space result and Slice 2. Marker genes were found by 'FindAllMarkers' function in Seurat. P value was calculated with the Wilcoxon Rank Sum test. **c**, Single-cell deconvolution of the bulk data. Top, t-SNE plot showed the clustering space of generated single cells and the rose chart showed the proportion of each predicted cell type. Bottom, distribution of predicted single cells. Different colors represented distinct cell types. **d**, The top, Histological annotation for stroma (red), melanoma (black), and lymphoid region (yellow). Bottom, single-cell decomposition result of each spot in the spatial reference data by Bulk2Space. **e**, The spatial abundance of different cell types in each spot in the tissue section was predicted by Bulk2Space. **f**, The expression correlation of the marker genes in six cell types between the Bulk2Space result and Slice 1. Marker genes were found by 'FindAllMarkers' function in Seurat. P value was calculated with the Wilcoxon Rank Sum test. **g**, The spatial expression of the marker genes in B cells, CAFs, macrophages, and malignant cells in predicted data, respectively. **h**, Spatial distribution of predicted B cells. Segmented areas, lymphoid tissue (purple), and melanoma region (black). **i**, Differentially expressed genes for lymphoid and melanoma tissue. **j**, Pathway enrichment analysis of the predicted B cells for lymphoid and melanoma tissue. *P* value was calculated with the accumulative hypergeometric distribution. Source data are provided as a Source Data file.

#### Bulk2Space integrates spatial gene expression and histomorphology in PDAC using discrete slices

##### *Spatial analysis of bulk PDAC by Bulk2Space*

Similarly, Bulk2Space was applied to reveal the spatial heterogeneity in bulk PDAC data using discrete slices of different PDAC tissues. Moncada[23] et al. performed scRNA-seq and ST[26] on PDAC tissues. As shown in Fig. S17a, the scRNA-seq data were used as a deconvolution reference, and one piece of a tissue section was used as a spatial reference (Slice 1). An additional slice (Slice 2) was used to synthesize bulk data. After deconvolution, single-cell transcriptomics data were generated from the bulk data. The expression of cell-type-specific marker genes between the generated single-cell profiles and Slice 2 were highly correlated (Fig. S17b). The generated single cells were then mapped to Slice 1, a spatial barcoding-based reference, by Bulk2Space (Fig. S17c).

To link the spatial distribution pattern of generated single cells with the histological feature of tissue regions, we investigated the molecular architecture of the annotated regions in Slice 1 (Fig. S17d). As shown, the spatial distribution of cancer, endothelial, and ductal cells predicted by Bulk2Space (Fig. S17e) was consistent with the histological annotation and the spatially resolved transcriptomics of cancer, interstitium, and duct epithelium regions. This was confirmed by the spatial expression of marker genes in these three histologically associated cell types (Fig. S17f). The gene expression correlation of all six cell types between the predicted spatially resolved single-cell transcriptomics data and Slice 1 was exhibited in Fig. S17g. The H&E staining of the slices were shown in Fig. S17h (Slice 1) and Fig. S17i (Slice 2).

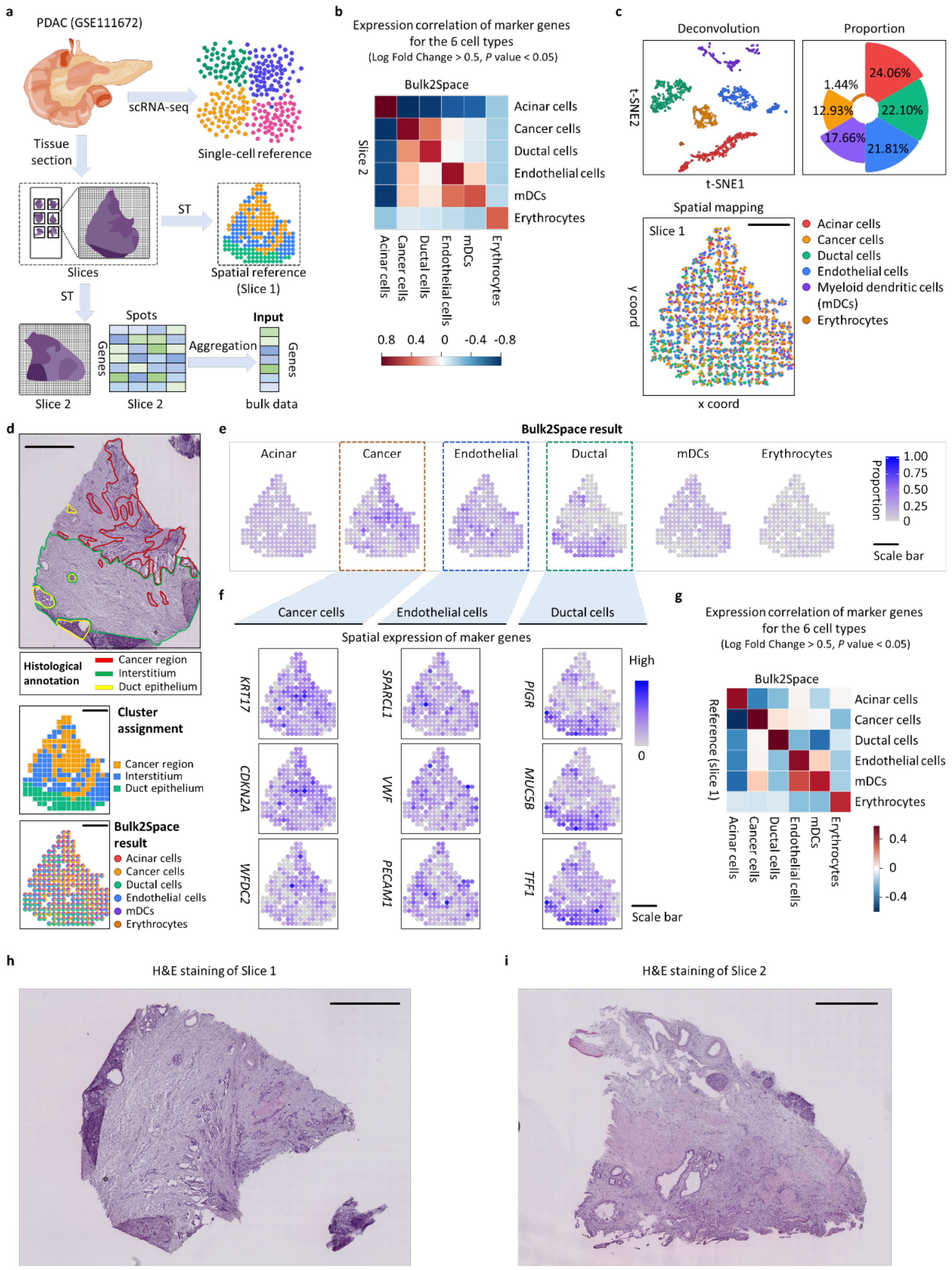

**Fig. S17**. **Spatially resolved analysis of PDAC by Bulk2Space using consecutive slices. a,** Synthesis of the bulk transcriptome data. The scRNA-seq data is used as the deconvolution reference, one slice of the sectioned tissue is employed as the spatial reference (Slice 1), and another slice (Slice 2) is used to synthesize the input bulk data. **b**, The expression correlation of the marker genes in six cell types between the Bulk2Space result and Slice 2. Marker genes were found by 'FindAllMarkers' function in Seurat. P value was calculated with the Wilcoxon Rank Sum test. **c**, Deconvolution of the bulk data into single cells. Top, t-SNE plot showed the clustering of generated single cells and the rose chart showed the proportion of each predicted cell type. Bottom, spatial distribution of predicted single cells. Scale bar, 1 mm. Different colors represented distinct cell types. **d**, Top, Histological annotation for cancer region (red), interstitium (green), and duct epithelium (yellow). Middle, Clusters of cancer region (yellow), interstitium (blue), and duct epithelium (green) from the spatially resolved trancsriptomics data. Bottom, Spatial deconvolution result of the bulk data by Bulk2Space. Scale bar, 1 mm. **e**, The spatial abundance of different cell types in each spot on the tissue section was predicted by Bulk2Space. Scale bar, 1 mm. **f**, The spatial expression of the marker genes in cancer (Left), endothelial (Middle), and ductal (Right) cells in predicted data. Scale bar, 1 mm. **g**, The expression correlation of the marker genes in six cell types between the Bulk2Space result and Slice 1. Marker genes were found by 'FindAllMarkers' function in Seurat. P value was calculated with the Wilcoxon Rank Sum test. **h**, H&E staining of Slice 1. Scale bar, 1 mm. **i**, H&E staining of Slice 2. Scale bar, 1 mm. Repeated experiments were not used to validate the results. Source data are provided as a Source Data file.

##### ***Cell types that exist in the reference dataset yet were predicted to be 0 by Bulk2Space***

The proportion of each cell type predicted by Bulk2Space was different from that in the reference scRNA-seq of PDAC. Notablely, four cell types in the reference data, including endocrine cells, monocytes, macrophages, and tuft cells, were not predicted in the generated single-cell data. Therefore, we further validated the distribution of the marker genes of these cell types. As shown in Fig. S18, the predicted distribution of marker gene expression in Slice 1 was consistent with the real spatial expression of these genes in spatially resolved transcriptomics data, which indicated that the prediction results of Bulk2Space were robust.

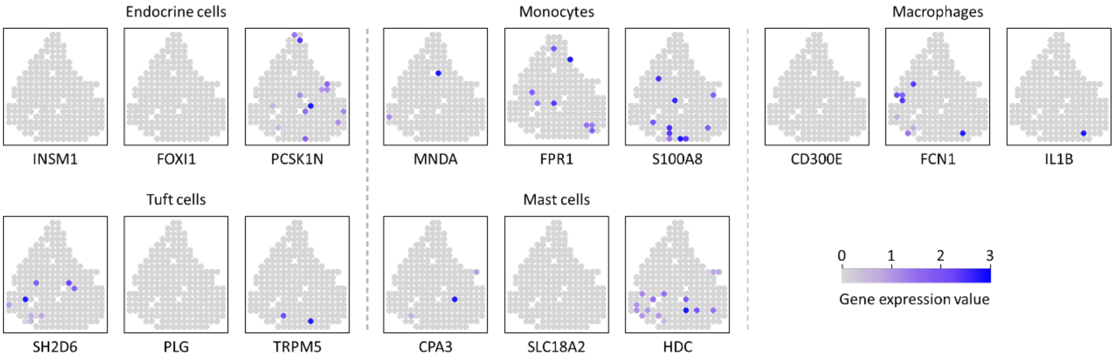

**Fig. S18**. **Spatial expression of cell-type-specific marker genes for cell types predicted absent in**

**the generated single-cell data by Bulk2Space**. The spatial expression of three marker genes for endocrine cells, monocytes, macrophages, tuft cells, and mast cells. Source data are provided as a Source Data file.

### Application of Bulk2Space in biological and clinical circumstances

#### Bulk2Space integrates spatial gene expression and histomorphology in PDAC using biological bulk data

In the validation of Bulk2Space's performance, consecutive slices were used as the spatial reference and synthetic bulk data, respectively. Here we discuss whether Bulk2Space could be applied to biological circumstance for spatial deconvolution of PDAC using biological bulk data.

As the supplementary information for Fig. 3 in the main text, results on real data had proved the potential application prospect of Bulk2Space. The clustering space of the generated single cells from pancreatic adjacent (PA) tissues was shown in Fig. S19a. The expression of marker genes for cell types between single-cell profiles generated from PA tissues by Bulk2Sapce and the single-cell reference data was highly correlated (Fig. S19b).

For pancreatic cancer tissues, spatial proportions of seven cell types predicted by Bulk2Space were shown in Fig. S19c. The spatial expression of cell-type-specific marker genes at spot (left) and single-cell (right) resolution for seven cell types, endothelial cells, fibroblasts, macrophages, myeloid dendritic cells (mDCs), Monocytes, plasmacytoid dendritic cells (pDCs), and red blood cells (RBCs) were illustrated in Fig. S19d. Compared with the spatial distribution of generated single cells, the spatial expression of cell-type-specific marker genes exhibited consistent patterns.

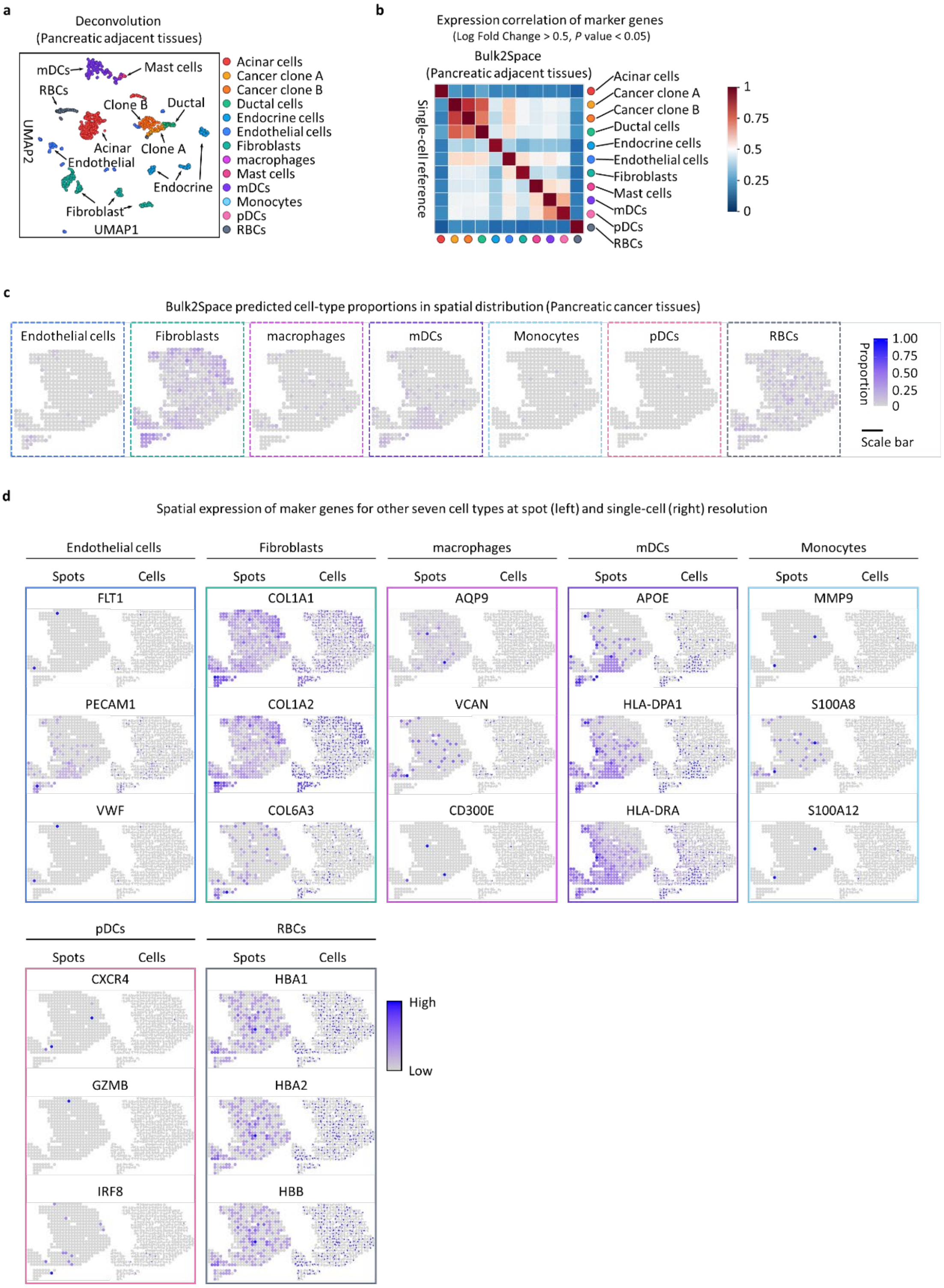

**Fig. S19**. **Spatially resolved analysis of the PDAC by Bulk2Space using biological data from pancreatic adjacent and cancer tissues**. **a**, Deconvolution results for bulk pancreatic adjacent tissues using Bulk2Space. **b**, Pairwise expression correlation of cell-type-specific marker genes between single cells generated by Bulk2Space and the single-cell reference for PA tissues. Marker genes were found by 'FindAllMarkers' function in Seurat. P value was calculated with the Wilcoxon Rank Sum test. **c**, Spatial distribution of the cell-type proportion predicted by Bulk2Space. **d**, Spatial expression of three marker genes for seven cell types at spot (Left) and cellular (Right) resolution. Source data are provided as a Source Data file.

#### Spatial deconvolution of biological bulk RNA-seq data derived from our in-house developed Spatial-seq technology

The Spatial-seq workflow is shown in Fig. S20a, the laser capture microdissection (LCM) was used to isolate brain regions and a spatial barcoding strategy was utilized for multiplexed RNA-seq. Specifically, the mouse brain was sliced into 14-µm sections from the coronal and sagittal directions. Each tissue slice was registered to a reference brain template provided by the Allen Brain Atlas (https://portal.brain-map.org/). After spatial registration, the anatomical regions of the mouse brain were delineated and annotated. The outlines, dissection sequences, and collectors of the brain regions were specified from the annotated data and then imported to an LCM instrument. LCM is a microscope-guided powerful cutting system incorporating UV light for contact- and contamination-free isolation of areas of interest from tissue sections[27]. Brain regions were dissected from the tissue by LCM according to the imported files and fell into the collector loaded with barcoded beads in advance. Subsequently, the tissue was lysed in the barcoded well, allowing mRNA to be captured by polyT tail on the surface of the magnetic beads. The captured mRNA was reversely transcribed to construct cDNA library and a paired-end sequencing was conducted to decode the spatial barcodes in the 3' end and detect RNA species in the 5' end of the cDNA.

In this study, 214 brain regions from three mice were sequenced following the Spatial-seq procedure. Two out of the 214 bulk RNA-seq data, the mouse isocortex and hypothalamus region, were used as two examples of Bulk2Space applications (Fig. S20b) as supplementary information to Fig. 4 and Fig. 5 in the main text.

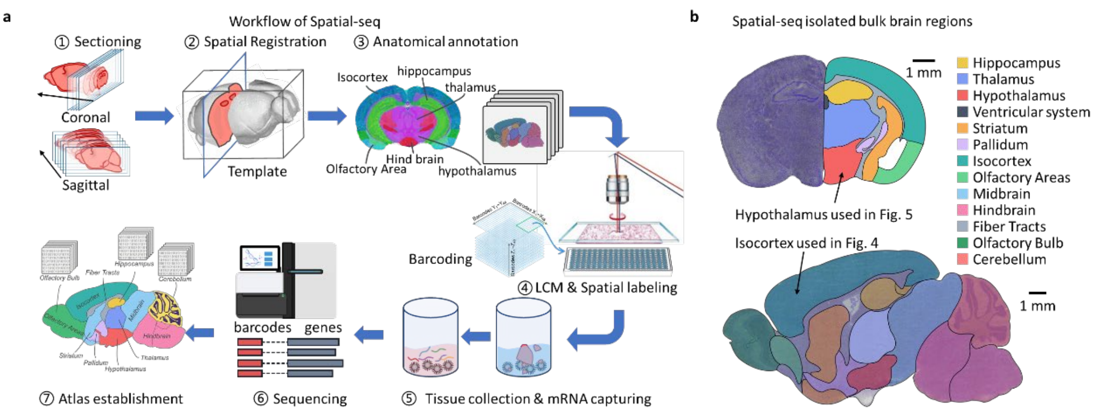

**Fig. S20**. **Workflow of Spatial-seq and two isolated regions of the mouse brain used in this study**. **a**, Spatial-seq workflow. The mouse brain was sectioned continuously from coronal and sagittal directions. For each slice, spatial registration was carried out to annotate each brain region to be isolated. Brain regions are dissected by LCM and labeled with a unique barcode and then followed with RNA-seq. **b**, Two isolated brain regions from two mice used in the main text. The hypothalamus region from a coronal section and the isocortex region from a sagittal section. The scale bar for each slice is shown in the figure. Source data are provided as a Source Data file.

#### Bulk2Space predicted the spatial expression of novel genes

As the complementary information to Fig. 5 in the main text, the spatial expression of novel marker genes of 9 cell types was shown in Fig. S21, with three marker genes for each cell type. Notably, these marker genes were not provided by the original MERFISH data.

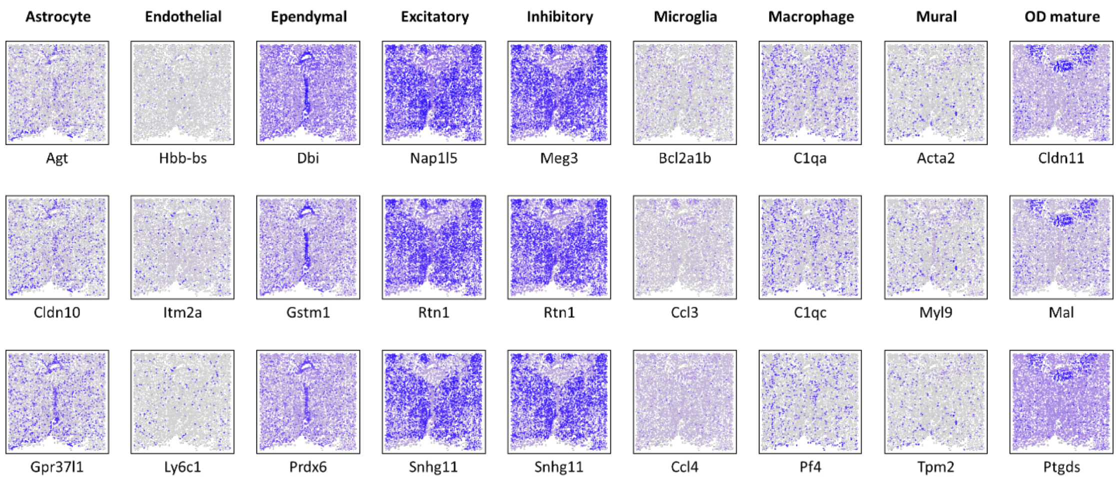

**Fig. S21. The spatial distribution of novel marker genes of each cell type predicted by Bulk2Space.** For each cell type, three marker genes absent in MERFISH data were predicted by Bulk2Space. Source data are provided as a Source Data file.

### Reference

1. Gierahn, T.M. et al. Seq-Well: portable, low-cost RNA sequencing of single cells at high throughput. *Nat Methods* **14**, 395-398 (2017).

2. Fan, X. et al. Spatial transcriptomic survey of human embryonic cerebral cortex by single-cell RNA-seq analysis. *Cell Res* **28**, 730-745 (2018).

3. Lake, B.B. et al. A single-nucleus RNA-sequencing pipeline to decipher the molecular anatomy and pathophysiology of human kidneys. *Nat Commun* **10**, 2832 (2019).

4. Aizarani, N. et al. A human liver cell atlas reveals heterogeneity and epithelial progenitors. *Nature* **572**, 199-204 (2019).

5. Vieira Braga, F.A. et al. A cellular census of human lungs identifies novel cell states in health and in asthma. *Nat Med* **25**, 1153-1163 (2019).

6. Zeisel, A. et al. Brain structure. Cell types in the mouse cortex and hippocampus revealed by single-cell RNA-seq. *Science* **347**, 1138-1142 (2015).

7. Wu, H., Kirita, Y., Donnelly, E.L. & Humphreys, B.D. Advantages of Single-Nucleus over Single-Cell RNA Sequencing of Adult Kidney: Rare Cell Types and Novel Cell States Revealed in Fibrosis. *J Am Soc Nephrol* **30**, 23-32 (2019).

8. Zilionis, R. et al. Single-Cell Transcriptomics of Human and Mouse Lung Cancers Reveals Conserved Myeloid Populations across Individuals and Species. *Immunity* **50**, 1317-1334 e1310 (2019).

9. Baron, M. et al. A Single-Cell Transcriptomic Map of the Human and Mouse Pancreas Reveals Inter- and Intra-cell Population Structure. *Cell Syst* **3**, 346-360 e344 (2016).

10. Green, C.D. et al. A Comprehensive Roadmap of Murine Spermatogenesis Defined by Single-Cell RNA-Seq. *Dev Cell* **46**, 651-667 e610 (2018).

11. Grun, D. et al. De Novo Prediction of Stem Cell Identity using Single-Cell Transcriptome Data. *Cell Stem Cell* **19**, 266-277 (2016).

12. Muraro, M.J. et al. A Single-Cell Transcriptome Atlas of the Human Pancreas. *Cell Syst* **3**, 385-394 e383 (2016).

13. Lawlor, N. et al. Single-cell transcriptomes identify human islet cell signatures and reveal cell-type-specific expression changes in type 2 diabetes. *Genome Res* **27**, 208-222 (2017).

14. Segerstolpe, A. et al. Single-Cell Transcriptome Profiling of Human Pancreatic Islets in Health and Type 2 Diabetes. *Cell Metab* **24**, 593-607 (2016).

15. Frishberg, A. et al. Cell composition analysis of bulk genomics using single-cell data. *Nat Methods* **16**, 327-332 (2019).

16. Newman, A.M. et al. Robust enumeration of cell subsets from tissue expression profiles. *Nat Methods* **12**, 453-457 (2015).

17. Chen, Z. et al. Inference of immune cell composition on the expression profiles of mouse tissue. *Sci*

*Rep* **7**, 40508 (2017).

18. Wang, J., Roeder, K. & Devlin, B. Bayesian estimation of cell type-specific gene expression with prior derived from single-cell data. *Genome Res* **31**, 1807-1818 (2021).
19. Satija, R., Farrell, J.A., Gennert, D., Schier, A.F. & Regev, A. Spatial reconstruction of single-cell gene expression data. *Nat Biotechnol* **33**, 495-502 (2015).
20. Xiong, X. et al. Landscape of Intercellular Crosstalk in Healthy and NASH Liver Revealed by Single-Cell Secretome Gene Analysis. *Mol Cell* **75**, 644-660 e645 (2019).
21. Stickels, R.R. et al. Highly sensitive spatial transcriptomics at near-cellular resolution with Slide-seqV2. *Nat Biotechnol* **39**, 313-319 (2021).
22. Saunders, A. et al. Molecular Diversity and Specializations among the Cells of the Adult Mouse Brain. *Cell* **174**, 1015-1030 e1016 (2018).
23. Moncada, R. et al. Integrating microarray-based spatial transcriptomics and single-cell RNA-seq reveals tissue architecture in pancreatic ductal adenocarcinomas. *Nat Biotechnol* **38**, 333-342 (2020).
24. Tirosh, I. et al. Dissecting the multicellular ecosystem of metastatic melanoma by single-cell RNA-seq. *Science* **352**, 189-196 (2016).
25. Thrane, K., Eriksson, H., Maaskola, J., Hansson, J. & Lundeberg, J. Spatially Resolved Transcriptomics Enables Dissection of Genetic Heterogeneity in Stage III Cutaneous Malignant Melanoma. *Cancer Res* **78**, 5970-5979 (2018).
26. Stahl, P.L. et al. Visualization and analysis of gene expression in tissue sections by spatial transcriptomics. *Science* **353**, 78-82 (2016).
27. Nichterwitz, S. et al. Laser capture microscopy coupled with Smart-seq2 for precise spatial transcriptomic profiling. *Nat Commun* **7**, 12139 (2016).
